# Resurrection of the island rule – human-driven extinctions have obscured a basic evolutionary pattern

**DOI:** 10.1101/025486

**Authors:** Søren Faurby, Jens-Christian Svenning

**Affiliations:** Section for Ecoinformatics & Biodiversity, Department of Bioscience, Aarhus University, Ny Munkegade 114, DK-8000 Aarhus C, Denmark; Department of Biogeography and Global Change, Museo Nacional de Ciencias Naturales, CSIC, Calle José Gutiérrez Abascal 2, Madrid 28006, Spain

**Keywords:** anthropocene, body size, evolution, islands, mammals

## Abstract

Islands are or have been occupied by unusual species, such as dwarf proboscideans and giant rodents. The discussion of the classical but controversial “island rule,” which states that mammalian body sizes converge on intermediate sizes on islands, has been stimulated by these unusual species. In this paper, we use an unprecedented global data set of the distributions and the body sizes of mammals and a novel analytical method to analyze body size evolution on islands; the analyses produced strong support for the island rule. Islands have suffered massive human-driven losses of species, and we found that the support for the island rule was substantially stronger when the many late-Quaternary extinct species were also considered (particularly, the tendency for dwarfing in large taxa). In this study, the decisive support generated for the island rule confirmed that evolution is markedly different on islands and that human impact may obscure even fundamental evolutionary patterns.

## Introduction

Before the arrival of humans, many oceanic islands housed bizarre mammal faunas. Dwarf proboscideans used to occur on Mediterranean islands, the Channel Islands in California, and the island of Timor in Southeast Asia, but all are extinct (Faurby and Svenning 2015). Similarly, giant rats were frequent on islands, with only a few species that are extant (Faurby and Svenning 2015), although in some cases with much reduced ranges, e.g., the Malagassy giant rat (*Hypogeomys antimena*) (Burney et al. 2008). In addition to these clades with numerous deviant island forms, many other clades also had a single or a few odd-sized island species, e.g., the extinct dwarf hippos of Crete and Madagascar and the extinct Sardinian giant pika (Stuenes 1989, Angelone et al. 2008).

These bizarre island mammals stimulated the proposal of the island rule, which states that mammalian body sizes converge on intermediate sizes on islands (Van Valen 1973). However, the island rule has been intensely debated in recent years and is viewed as both a near universal rule (Lomolino et al. 2011) and a sample or publication artifact (Meiri et al. 2008, Raia et al. 2010), with intermediate positions also argued (Welch 2009). Both the opponents and the proponents of the island rule acknowledge the apparent abundance of giants and dwarfs on islands (Meiri et al. 2008, Lomolino et al. 2011). The two schools have strongly argued whether the island rule represents a general evolutionary pattern, the idiosyncratic changes in individual lineages or even the human tendency to see patterns in all datasets (Van Valen 1973, Meiri et al. 2008, Raia et al. 2010, Lomolino et al. 2011).

Critics of the island rule argue two primary points, both of which we overcome in the present study. The first point concerns sampling bias. The studies that support the island rule have generally been meta-analyses of published comparisons between the mainland and island populations of the same species (Van Valen 1973, Lomolino et al. 2011). As discussed for the related Bergmann’s rule (Meiri et al. 2004), these studies may be a nonrandom subset of all populations and therefore a significant pattern matching expectation may be generated by a reporting bias. In this study, we removed the possibility for such sampling bias by generating and analyzing a database that contained the body sizes for approximately 99% of all extant and recently extinct species of mammals (see Materials and Methods).

The second critique of the basis for the island rule is that of phylogenetic nonindependence, because previous studies showed diminished support for the rule when the phylogeny was accounted for in the intraspecific analyses (Meiri et al. 2008, McClain et al. 2013). This problem is a form of pseudo-replication that inflates the estimates of precision and thereby potentially generates false significances. The magnitude (or existence) of this problem, however, depends on what model is used as the null model. The classical studies, which compared only sister lineages (e.g., Lomolino 1985), are compatible with body sizes that evolved via simple models such as the Brownian motion (Felsenstein 1985) or the Ornstein-Uhlenbeck (OU) models (Hansen 1997). The studies that analyzed the ratios between the sizes of island and mainland mammals in a phylogenetic context (e.g., Meiri et al. 2008) might also be compatible with such models, when one assumes an identical age for all island populations or that the island populations have reached a new equilibrium size.

However, with the assumption that the rate of evolutionary change is a function of traits, which are also evolving, i.e., via the correlation between generation length and evolutionary rate (Welch et al. 2008, Thomas et al. 2010), phylogenetic nonindependence is a problem for studies that do not integrate phylogeny. Such a correlation is not a problem for studies that incorporate phylogeny and that focus on the ratios between the island and mainland species, but these studies also require an identical age of all island populations or that the island populations have reached a new equilibrium size. Imagine, for example, an analysis that contained two sets of rodent mainland /island sister pairs with short generation times and therefore potentially fast evolutionary rates and that contained two sets of elephant mainland/island species pairs with longer generation times and therefore potentially lower evolutionary rates. If the rate is evolving over time, the comparisons of the magnitude of change between the species pairs will need phylogenetic correction because larger relative differences between the mice species pairs than the elephant species pairs could be a null expectation. Irrespective of whether the rate is evolving, however, the null expectations would remain a 50% decrease in size in both the mice and the elephants. To solve the potential problem of phylogenetic nonindependence without requiring an identical age of all island populations or that the island populations have previously reached a new equilibrium size, we restricted the analyses to focus only on the directionality and not on the magnitude of change (see the Materials and methods). We stress that this restriction did not imply that body size did not evolve as a Brownian motion process (there are strong indications that it often does (Blomberg et al. 2003)) but that our analysis (explained below) did not make that assumption. Moreover, the analysis was almost independent of the assumed model of body size evolution.

In addition to the potential problems with the studies that support the island rule, the primary interspecific study that dismisses the island rule (Raia et al. 2010) also has potential problems. The study used a somewhat incomplete body size database (Smith et al. 2003) and a partially outdated phylogeny (Bininda-Emonds et al. 2007). Raia et al. (2010) also included bats, whereas the classic studies that support the island rule focused on non-flying mammals (Lomolino 1985) or analyzed bats separately (Lomolino 2005). If supported, the island rule is likely a consequence of island isolation, and the substantially lower levels of endemism in bats than in non-flying mammals (Weyeneth et al. 2011) indicates that the island bat fauna is less isolated compared to non-flying mammalian fauna. Thus, the island rule would be expected to establish a weaker pattern for bats than for non-bats.

In this paper, we reanalyzed the magnitude of the island rule in an interspecific context using a novel, near-complete body size database and a recent mammalian phylogeny (Faurby and Svenning 2015) solely focusing on the directionality and not on the magnitude of body size changes in island lineages. To determine the potential importance of the factors responsible for the apparent lack of support for the island rule in the earlier studies that integrated phylogenetic relationships between species, we estimated the effects of including or excluding bats and extinct species and different definitions of islands.

## Materials and methods

### Data generation

For all analyses, we used the taxonomy and the phylogeny of a recent mammalian phylogeny, which included all species with dated occurrences within the last 130,000 years, but no likely chronospecies (Faurby and Svenning 2015). Notably, most extant mammal species existed throughout this period and therefore coexisted with the extinct species, and there is increasing evidence that Homo sapiens were the primary cause of these extinctions (Sandom et al. 2014), particularly on islands (Turvey and Fritz 2011).

We generated a new body size database, which included almost all species of mammals (5673 of 5747 species; of the 74 species without data, 8 represented extinct, but undescribed, species). The new database was partly based on an older database (Smith et al. 2003) but was heavily modified. The information for 3629 of the 5673 species was used from the older database, but our complete database contained information from a total of 709 separate data sources (644 articles published in 146 separate journals, 55 books, 8 web resources and personal information from 2 experts; the complete database is available in the Supplementary Data, in addition to information on which islands all island endemic species are found). For the species for which the weight data were not available, the weights were generally estimated with the assumption of strict isometries for related similar sized species. The isometry was generally assumed for forearm length in bats and body length (excluding tail) for the remaining species, but other measures were also used occasionally.

We scored island endemic or remainder as a binary character and defined island endemics based on three definitions. The loose and classical definition was any species endemic to any area, which are the oceanic islands at the current sea levels. The species that are currently restricted to islands (or were restricted until their extinction in historical times) but with former Holocene occurrences on the mainland, e.g., the Tasmanian devil (*Sarcophilus harrisii*) and the Tasmanian tiger (*Thylacinus cynocephalus*) (Johnson and Wroe 2003), were not scored as island endemics. The strict definition was for any species that was not found on any continent or any island connected to a continent during the ice ages (i.e., any island for which the deepest water-level between the island and a continent was less than 110 meters deep). Using this restricted definition, the island endemics were species for which the majority of their evolutionary history were restricted to islands instead of species that happened to be on islands with the current sea levels. For the few species that evolved by rapid speciation since the last ice age on land-bridge islands (e.g., Anderson and Hadley 2001), this definition may be overly restrictive because the species would have been island endemics for their entire evolutionary history. Therefore, we also used a semi-strict definition, which was a relaxation of the strict island definition, and any species that did not occur on large land-bridge islands (larger than 1000 km^2^) were also scored as island endemics. We acknowledge that this threshold was somewhat arbitrary, but rapid speciation since the end of the last ice age likely required a small population size and therefore a limited area. The largest island with a strong candidate for such recent speciation would be Coiba (503 km^2^) with the endemic agouti *Dasyprocta coibae*, whereas the colobus monkey *Procolobus kirkii* from Zanzibar (1658 km^2^, the smallest land-bridge island above 1000 km^2^ that contained an endemic species) appeared to have been isolated for substantially longer than the end of the last ice age (Ting 2008).

### Analyses

The phylogeny used in this study consisted of the 1000 separate, random fully bifurcating trees from a posterior distribution of trees that represented the phylogenetic uncertainty from Faurby and Svenning (2015). Separate analyses were initially performed for each of the 1000 trees after which the results from each tree were combined.

For each island endemic species (IE), we found the largest clade that contained only island endemics (C_Island_) and the smallest clade that contained both island endemics and nonendemics and removed all members of C_Island_ from this clade (hereafter C_Mainland_). We then estimated ancestral log_10_(Weight) for all C_Island_ and C_Mainland_, assuming Brownian motion. With the removal of the island endemics for the calculation of C_Mainland_, we allowed body size evolution to differ between island and mainland clades but did not enforce such differences. Following this procedure, we sampled all the island endemic species in random order, listed all members of C_Island_ and, if the sampled species was not a member of the C_Island_ of any previously sampled species, noted the size of the (Size_Mainland_) and whether this size was smaller or larger than Size_Island_. Therefore, our end products were a vector of ancestral mainland weights for independent island invasions and a corresponding vector with binary information on whether the island invaders were smaller than the mainland ancestors. To reduce the effects of measurement errors on weight, we discarded from further analyses all island invasions for which the difference in weight between Size_Mainland_ and Size_Island_ was smaller than 10%. Supplementary analyses were performed using 0%, 5%, 15% and 20% weight difference thresholds, but the results changed very little, although there was a tendency for a weaker island rule with the 0% threshold, which was likely a consequence of the increased noise in the data (see Supplementary Figures S1-S5)

We then fitted zero to the 4^th^ degree polynomial models of the probability of size decrease as a function of the Size_Mainland_ using a logistic regression 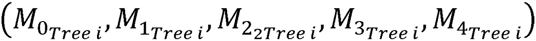 and calculated their respective AIC weights 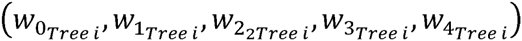. For all the potential values of Size_Mainland_ between 0.0 and 6.0 (i.e., untransformed weights between 1 g and 1 ton) in steps of 0.1 for all models, we then calculated the means and the variances, 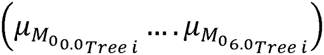 and 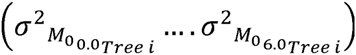, respectively, for the untransformed fitted values for all five models.

The results were thereafter combined for all five models for each potential value of C_Mainland_ as a mixture of the normal distributions from the five models, with the weight of each model equal to the AIC weight. Therefore, the combined result was that the predicted effect for any k would be in the distribution 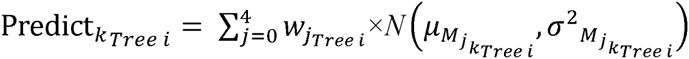. Following this procedure, the results were combined for all trees as 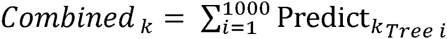. Finally, the median and several quantiles for the *Combined* _k_ were transformed into probabilities.

We tested the effect of the definition of an island endemic, the potential effect of the anthropogenic extinctions to bias the results and the effect of including bats in the analysis. The analysis was performed separately for each of the twelve combinations of the three definitions of island endemics (classical, semi-strict, strict), for the exclusion or inclusion of bats and for the exclusion or inclusion of extinct species.

All analyses were performed in R 3.0.2 (R Core Team. 2013) using functions from the libraries ape (Paradis et al. 2004), phylobase (R Hackathon et al. 2014) and qpcR (Spiess 2014).

## Model justification

Because our analysis included approximately 99% of all mammal species, the issue of publication bias was dismissed. However, we acknowledge that small biases might remain because of taxonomic practices, e.g., whether island populations that diverged more in size from their mainland relatives were more likely to be classified as separate species. One example of such a small bias is the island endemic pygmy sloth (*Bradypus pygmaeus*) (Anderson & Handley, 2001). These questionable populations or species are generally found on land-bridge islands (as with the pygmy sloth), and therefore, this type of bias is a problem that should affect only the classical or semi-strict but not the strict island endemic definition.

However, a problem noted by Meiri et al. (2008) might have affected our analysis. For the two vectors A and B, the correlation between B and B/A is significantly negative the majority of the time when A and B are independent. Meiri et al. (2008) extended this basic mathematical result to the island rule and argued that the island rule pattern would occur when the body sizes of the populations or species on the islands were independent of the body sizes of their mainland relatives. To assess the effect of this relationship, we randomized the body sizes of all species, of all species within families, or of all species within genera for the analysis with the apparent strongest island rule (i.e., strict island endemic definition, excluding bats and including extinct species). For this analysis, the body sizes were randomized anew for each of the 1000 trees.

## Results

Strong support for the island rule was provided when bats were excluded from the analysis but only weak support when the bats were included. Among the 12 combinations of island-type definitions and included species, the strongest support for the island rule (measured as the difference between the predicted probability for size increase species for species with a size of 1 ton and 1 gram) was with the strict island definition and the exclusion of bats but the inclusion of extinct species (Table 1, Figure 1, Figures S1-S5). The inclusion of bats in the analysis consistently led to markedly lower support for the island rule, and the addition of the bats removed or at least reduced the tendency for small mammals to increase in size on islands. The inclusion of the extinct species and the application of the strict or semi-strict island definitions provided stronger support for the island rule, but only when bats were excluded.

**Figure 1.**
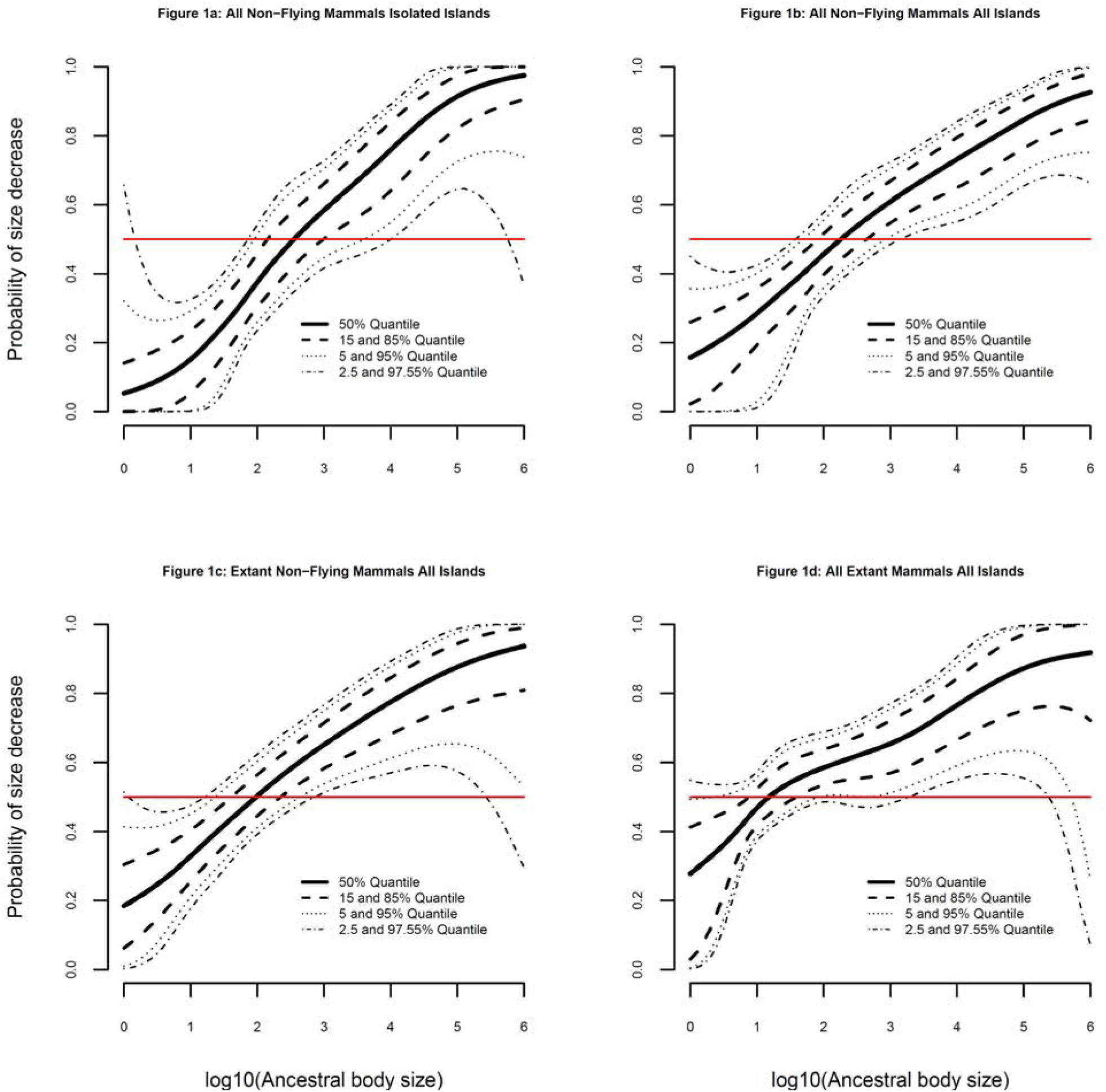
Relationships between ancestral body size and directionality of evolutionary size change after island invasion. The estimated probability of a size decrease as a function of the ancestral body size of the island invading clade. The thick black line shows the median of the distribution of potential predicted values, whereas the three stippled lines show the 2.5/97.5%, 5/95% and 15/85% quantiles. Because the response variable is binary, the values below the horizontal line indicate that a clade is most likely to increase in size, and the values above the horizontal line indicate that a clade is most likely to decrease in size. The first panel shows the relationship for non-flying mammals, including both extinct and extant species for isolated islands. The last panel shows the relationship for both flying and non-flying mammals but only for extant species and using all islands. The three differences between the panels are changed one by one; the last three panels use all the islands, the last two panels only analyze extant species and the last panel analyzes all mammals without restricting the analysis to non-flying species.

**Table 1.** The estimated strength of the island rule under the 12 different analyzed scenarios

| **Island definition** | **Including extinct species** | **Including bats** | **log10 of the body size with equal probability of size increase and decrease** | **Difference between the predicted probability of size increase for species of 1 ton and 1 gram** |
| --- | --- | --- | --- | --- |
| **Classical** | **No** | **Yes** | 1.3 | 0.641 |
| **Classical** | **No** | **No** | 2.1 | 0.753 |
| **Classical** | **Yes** | **Yes** | 1.6 | 0.602 |
| **Classical** | **Yes** | **No** | 2.4 | 0.769 |
| **Semi-strict** | **No** | **Yes** | 1.5 | 0.578 |
| **Semi-strict** | **No** | **No** | 2.4 | 0.867 |
| **Semi-strict** | **Yes** | **Yes** | 2.0 | 0.583 |
| **Semi-strict** | **Yes** | **No** | 2.6 | 0.893 |
| **Strict** | **No** | **Yes** | 1.4 | 0.589 |
| **Strict** | **No** | **No** | 2.4 | 0.893 |
| **Strict** | **Yes** | **Yes** | 2.0 | 0.616 |
| **Strict** | **Yes** | **No** | 2.7 | 0.922 |

The definitions of island endemics and the exclusion or inclusion of bats and extinct species also changed the shape of the relationship between body size and body size change on islands, in addition to influencing the magnitude of the island rule. For the strict island definition, when bats were excluded but extinct species were included, the apparent optimal size (the body size for which size increases and decreases are equally likely) was 500 gram (10^2.7^ g). On the other hand, for the classical island definition, when bats were included and, extinct species were not included, the optimal size was only 20 gram. (Table 1).

Our analysis of the effect of randomization of body sizes showed no support for the island rule under realistic randomization scenarios. Of the average of 143.3 independent island invasions from each of the 1000 separate trees, an average of 38.5 consisted of island endemic genera. When these genera were removed from the analysis with randomization within genera, almost no relationship between the body size and the directionality of size change was detected (Figure 2a). A slightly stronger but still weak pattern was found when the island endemic genera were included in the analysis (Figure 2b); however, randomization at the family level (Figure 2c) was required to falsely generate a pattern with substantial support for the island rule (the pattern with complete randomization was almost identical to the pattern for randomization at the family level, results not shown). The results of these randomizations strongly suggested that the support for the island rule was not an analytical artifact.

**Figure 2.**
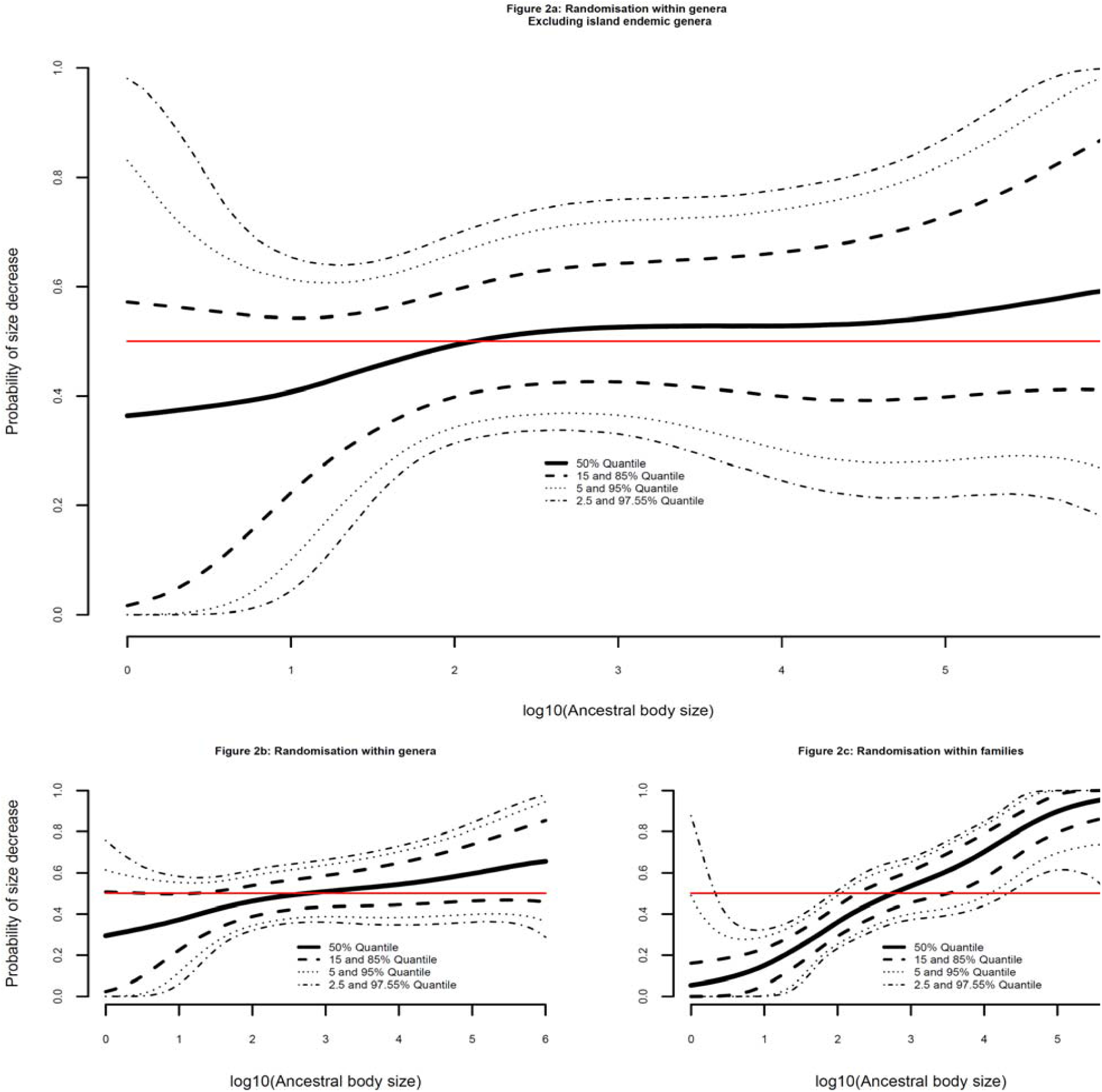
Relationships between ancestral body size and directionality of evolutionary size change for randomized body sizes. All panels show the effect of randomization of body sizes for the strict island rule, when extinct species are included but bats are excluded (analogous to Figure 1d). In Figure 2a, body sizes are randomized across all genera but with all island endemic genera excluded from the analysis, whereas in Figure 2b, body sizes are randomized across all species within genera and the island endemic genera are included in the analysis. In Figure 2c, body sizes are randomized across all species within families.

## Discussion

The validity of the island rule was clearly supported with our results. Therefore, we suggest that part of the explanation for the lack of evidence for the island rule in the earlier interspecific study (Raia *et al.* 2010) was not because of the incorporation of the phylogeny, as the authors suggested, but because of the choice to disregard the ecological difference between flying and non-flying mammals and, to a lesser extent, the choice of definition of island endemics and the incomplete inclusion of historically extant species (i.e., species that occurred within the Late Pleistocene or Holocene). In this regard, we do not state explicitly that the models that excluded bats in our analysis provided a better fit to the data than the models that included bats; however, we do state that the estimated effect of body size (i.e., the island rule) is substantially stronger in the models that excluded bats.

The effects of including or excluding land-bridge islands and bats into the analysis can potentially be seen as two sides of the same story. If the primary factor for the island rule was ecological release, the rule would only be realized on islands with reduced numbers of predators or competitor species. Therefore, the island rule would not apply or would be much less applicable to the land bridge islands, which were part of the continental mainland during the last glaciation, in addition to many earlier periods during the Pleistocene. The species on the land-bridge islands would have experienced similar faunas as on the current mainland for a large part of their evolutionary history, and therefore these species would not have experienced ecological release, or only relatively brief release. Similarly, island bats were not likely to experience a significant ecological release because the primary predators of bats are birds such as raptors and owls (Rydell and Speakman 1995). These birds are typically strong fliers and therefore even long isolated islands tend to harbor well-developed predatory bird faunas. Thus, for the native bat fauna on land-bridge islands, predator release would be limited or would not occur.

Our focus on interspecific patterns enables us to disentangle the different factors driving the island rule. The selection of immigrants for larger body sizes (as discussed in Lomolino, 2011) could potentially be important for relatively new intraspecific comparisons. Considering the evolutionary rates over small to medium time scales (Gingerich 2001), any effect of immigrant selection would disappear in interspecific analyses unless other selective forces were maintaining the changed body size (see Jaffe et al. (2011) for a similar argument regarding body size evolution in island tortoises). Therefore, our results indicated that selection caused by the novel ecology on islands was driving both the dwarfing and gigantism observed in different lineages.

With the arrival of humans, island faunas suffered severe extinctions. Our data set included 589 non-flying and 223 flying island endemics based on the strict island definition, with overall late-Quaternary extinction rates of 20% and 4%, respectively (the corresponding number for all the islands was 916 non-flying and 323 flying species with extinction rates of 13% and 3%, respectively; see Supplementary Data). These extinctions are often tightly linked to human arrival and to evidence of human hunting or other anthropogenic factors (Turvey and Fritz 2011). Based on our analysis, the inclusion of the extinct species strongly increased the support for the island rule. The incorporation of the extinct island species was previously advocated for ecological studies (Griffiths et al. 2009, Hansen and Galetti 2009), but our results highlighted the necessity to also include these species in evolutionary studies. The recent human-driven extinctions most likely obscured signals related to the long-term evolutionary responses to island environments, for example, the elimination of the most specialized of the island lineages (Lomolino et al. 2013). In this regard, we note that we expect both dwarfing and gigantism to be an evolutionary consequence of predator release. Therefore, the species that showed the largest changes in body size on islands would be expected to be the most sensitive to predation by humans or by our commensal animals.

The apparent optimal body size of 500 gram determined in our analysis using the strict island model and excluding bats but including extinct species (the model that showed the strongest support for the island rule) was similar to an estimate of optimal body size derived from the patterns in the intraspecific changes for terrestrial mammals on islands, which was 474 grams (Lomolino, 2005). However, several arguments against a global optimal body size have been developed (discussed in Raia et al. (2010)), and the similarity of the above results was possibly accidental. The potential accidental nature of the similarity of these results was also supported by the variation in the suggested global optimal size, if such an optimal size can be determined, with estimates of both 100 gram and 1 kg suggested previously (Brown et al., 1993; Damuth, 1993).

In this study, the decisive support for the island rule highlighted that the function of island ecosystems is fundamentally different from that of mainland systems (cf. Millen 2006) and that these differences drive divergent evolutionary dynamics on islands and the mainland. Notably, our results were consistent with the weakening of ecological interactions on islands that caused body sizes to shift to intermediate biomasses, irrespective of the ancestral body size or the phylogenetic lineage. Conversely, the strong support for the island rule also implied that much of the large variation in body sizes or the repeated evolution of similar maximum body sizes in mainland systems (Smith et al. 2010) was a consequence of the intense ecological interactions in these settings.

## Supplementary Materials

Figures S1-S5.

Supplementary data. A database of body size and island species endemicity.

**Figure S1.**
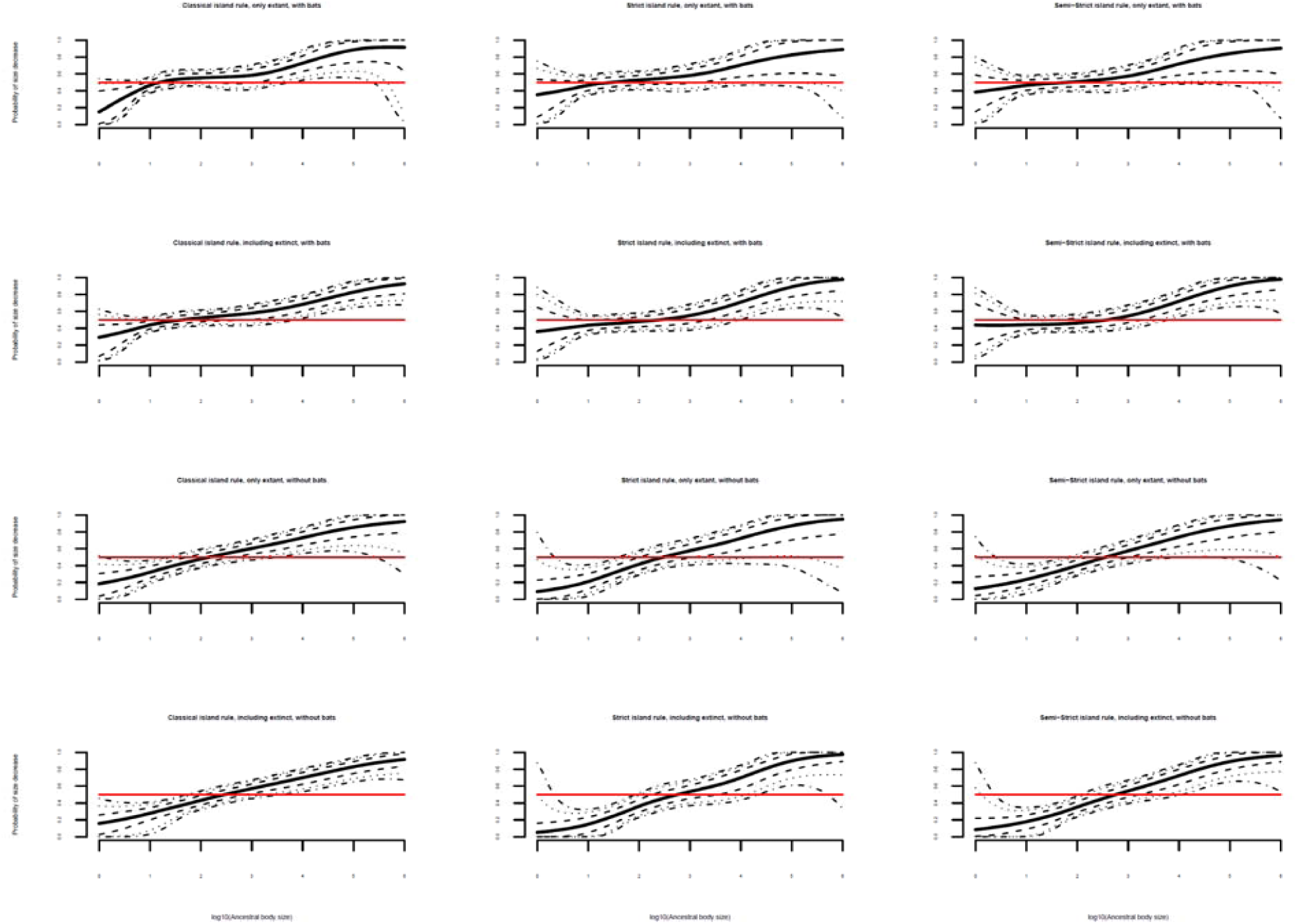
Relationships between ancestral body size and directionality of evolutionary size change after island invasion for the 12 separate analyses without any threshold for a minimum size difference between island and mainland clades. The structure and meaning of the individual lines in each panel are identical to those in Figure 1.

**Figure S2.**
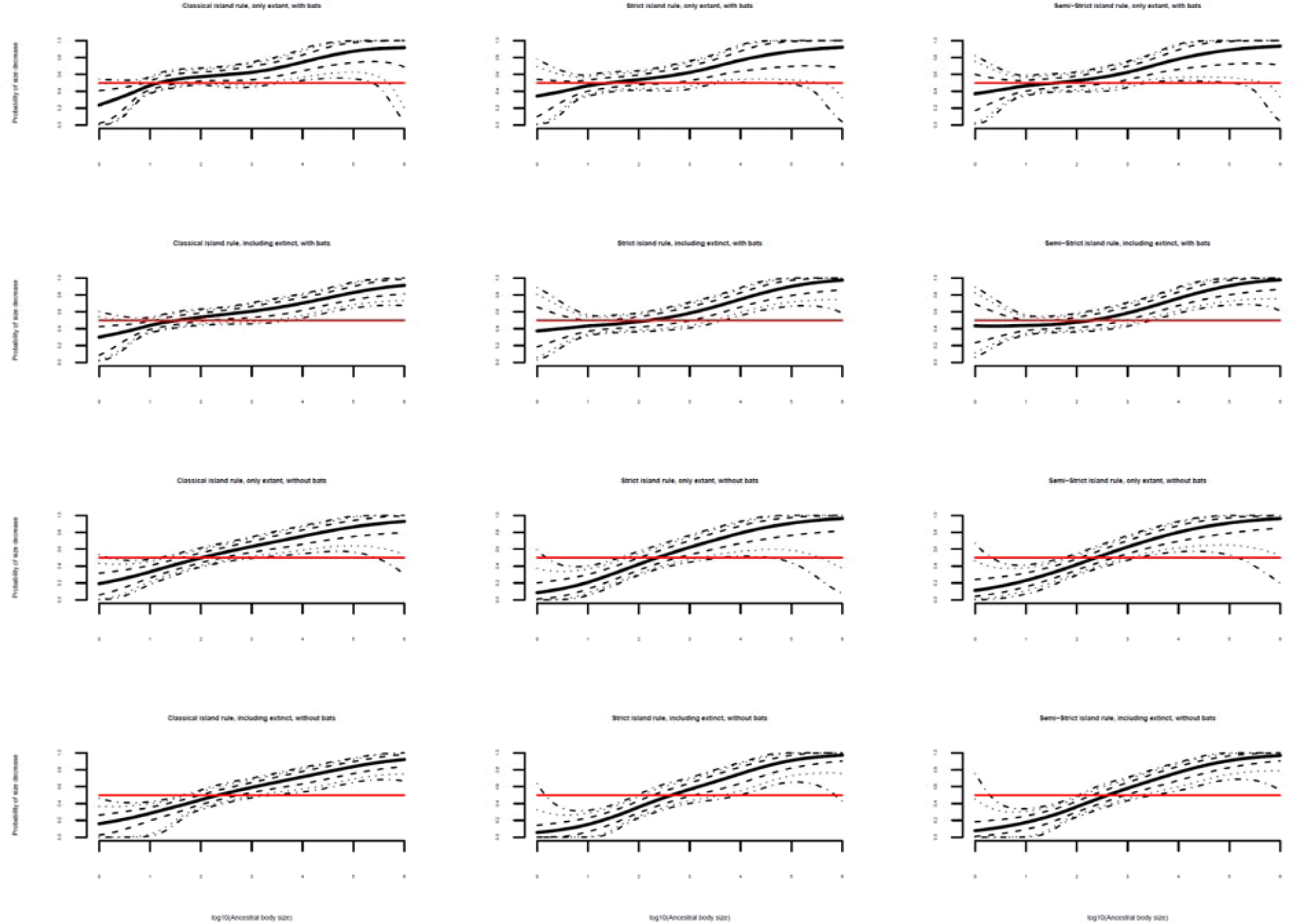
Relationships between ancestral body size and directionality of evolutionary size change after island invasion for the 12 separate analyses with a threshold for a minimum size difference between island and mainland clades of 5%. The structure and meaning of the individual lines in each panel are identical to those in Figure 1.

**Figure S3.**
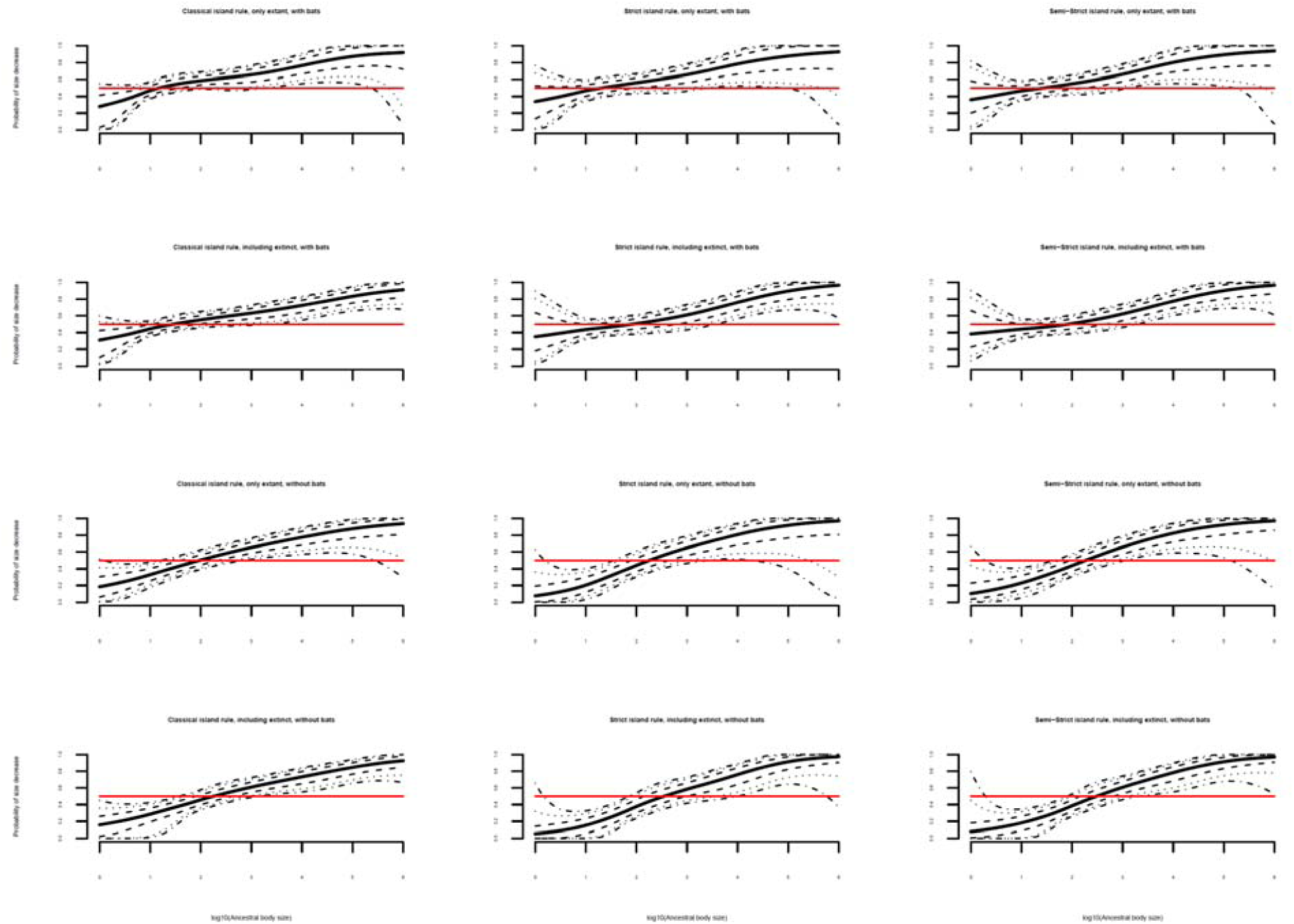
Relationships between ancestral body size and directionality of evolutionary size change after island invasion for the 12 separate analyses with a threshold for a minimum size difference between island and mainland clades of 10%. The structure and meaning of the individual lines in each panel are identical to those in Figure 1.

**Figure S4.**
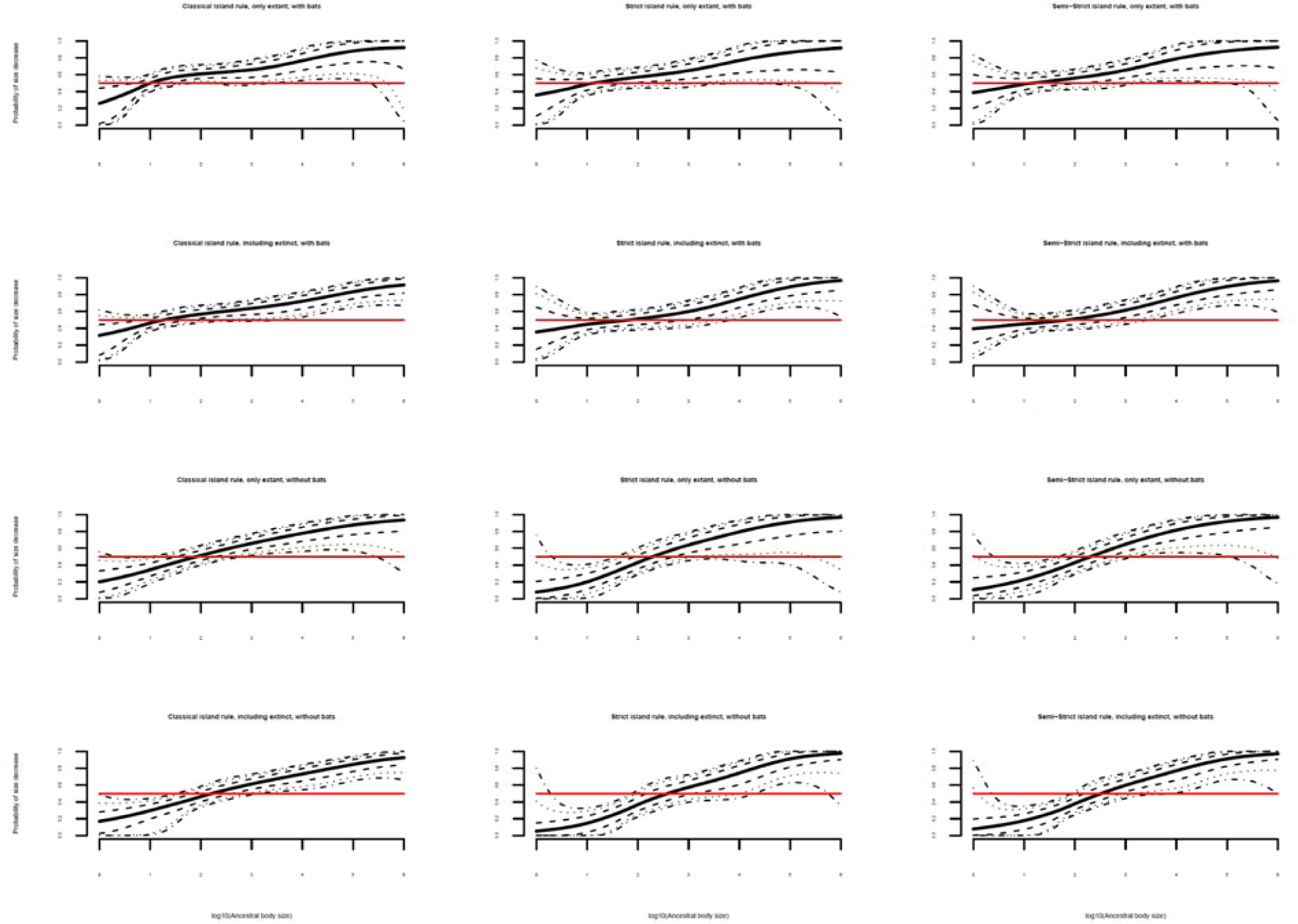
Relationships between ancestral body size and directionality of evolutionary size change after island invasion for the 12 separate analyses with a threshold for a minimum size difference between island and mainland clades of 15%. The structure and meaning of the individual lines in each panel are identical to those in Figure 1.

**Figure S5.**
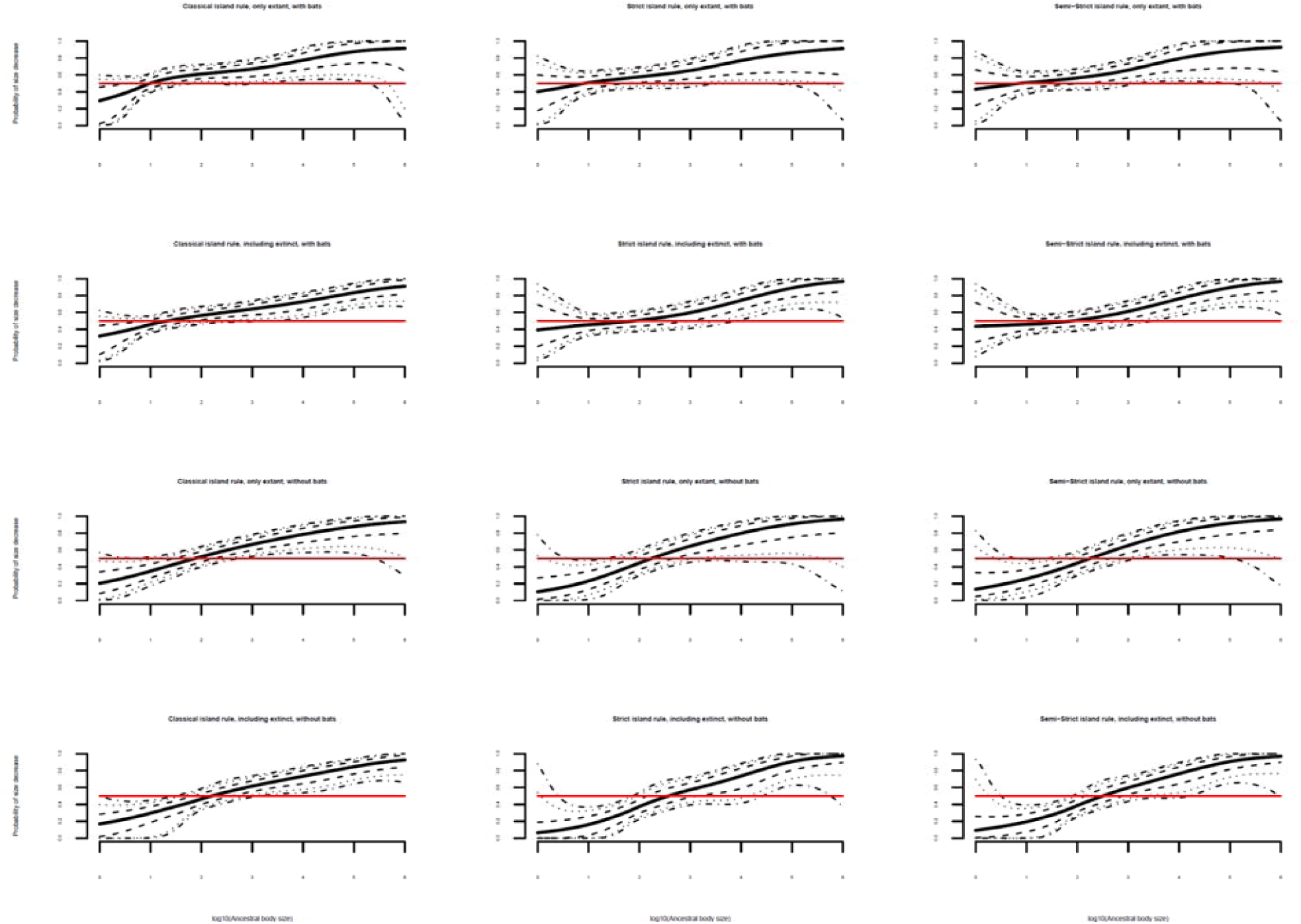
Relationships between ancestral body size and directionality of evolutionary size change after island invasion for the 12 separate analyses with a threshold for a minimum size difference between island and mainland clades of 20%. The structure and meaning of the individual lines in each panel are identical to those in Figure 1.

